# Spatial selective auditory attention is preserved in older age but is degraded by peripheral hearing loss

**DOI:** 10.1101/2024.05.30.596568

**Authors:** Andrea Caso, Timothy D Griffiths, Emma Holmes

## Abstract

Interest in how ageing affects attention is long-standing, although interactions between sensory and attentional processing in older age have not been systematically studied. Here, we examined interactions between peripheral hearing and selective attention in a spatialised cocktail party listening paradigm, in which three talkers spoke different sentences simultaneously and participants were asked to report the sentence spoken by a talker at a particular location. By comparing a sample of older (N = 61; age = 55– 80 years) and younger (N = 58; age = 18–35 years) adults, we show that, as a group, older adults benefit as much as younger adults from preparatory spatial attention. Although, for older adults, this benefit significantly reduces with greater age-related hearing loss. These results demonstrate that older adults without hearing loss retain the ability to direct spatial selective attention, but this ability deteriorates with age-related hearing loss. Thus, reductions in spatial selective attention likely contribute to difficulties communicating in social settings for older adults with age-related hearing loss. Overall, these findings demonstrate a relationship between mild perceptual decline and attention in older age.

## Introduction

How cognitive functions change with healthy ageing is a question of long-standing interest. Some cognitive abilities (e.g., working memory, visuo-spatial ability) appear to deteriorate with older age, whereas others (e.g., crystallised intelligence) improve^1^. Previous studies have examined visual and auditory spatial attention in older listeners, which is critical for environmental and social interactions. However, it is challenging to isolate changes in attention from the sensory processing of attended stimuli due to declines in hearing or vision.

Some studies have claimed that age-related changes previously attributed to attention are not present after controlling for sensory processing^2–4^, which is usually achieved by allowing older adults greater time to utilise an attentional cue or by equating baseline performance. However, these methods implicitly assume that changes in sensory processing are independent of attention, and do not consider possible interactions between top-down and bottom-up processes^5^. Yet, age-related declines in sensory processing may affect the allocation of attention—by degrading the sensory representations that people utilise to orient attention or, conversely, through increased reliance on attentional processes to compensate for degraded sensory processing. Crucially, such sensory–cognitive interactions could explain why previous studies on attentional changes in older age have found conflicting results^6^. In addition, sensory declines are extremely prevalent, so sensory–cognitive interactions are necessary for understanding cognitive processes in the vast majority of the ageing population, and could also provide insights into recently-established links between degraded peripheral function and risk of dementia^7,8^. Here, we assessed the relative contributions of age-related changes in attention that are dependent on peripheral processing, and those that are independent of peripheral processing, by explicitly modelling sensory–cognitive interactions.

We assessed sensory–cognitive interactions in a cocktail-party listening paradigm, in which participants hear sentences spoken by multiple talkers and are cued to selectively attend to a sentence at a particular location. This task involves endogenous attentional orienting (a type of ‘covert’ attention^9^). Crucially, hearing loss affects approximately 40% of people aged 55 and above^10^, and difficulties understanding speech in the presence of competing speech are not fully explained by measures of peripheral function, such as pure-tone audiometry^11,12^. Thus, changes to endogenous attention as a result of age, hearing loss, or both could help to explain why these situations are particularly difficult for older adults.

For young adults with normal hearing, directing spatial attention improves speech intelligibility in cocktail-party paradigms^13,14^, which appears to be underpinned by preparatory activity in fronto-parietal regions of cortex during the cue-target interval^15,16^ that may facilitate performance^17^. However, people with sensorineural hearing loss may not utilise spatial attention to the same extent. For example, Best *et al.*^18^ found that young adults (19–42 years) with hearing loss gained a significantly smaller improvement in intelligibility from a visual cue that informed them about the location of an upcoming target talker, compared to young adults with normal hearing. In addition, Holmes *et al.*^19^ found that EEG responses elicited by a visual cue for attention were weaker in children (aged 7–16 years) with than those without early-onset sensorineural hearing loss. Crucially, these studies of *preparatory* spatial attention allow auditory attention to be measured without confounding differences in peripheral auditory processing between people with and without hearing loss.

Yet, given the aforementioned studies tested young participants, we do not understand how *age-related* hearing loss affects preparatory spatial attention. On one hand, poorer spatial acuity may reduce the ability to orient spatial attention: Hearing loss degrades spectrotemporal resolution^20–22^, which can affect binaural and monaural spatial cues^23^. This could cause difficulty orienting selective attention to filter out distracting sounds, effectively broadening the focus of spatial attention^24^ and impairing sound segregation. Given that age-related hearing loss increases the minimal audible angle for discriminating spatial locations (to about 4 degrees, compared to approximately 1 degree in people without hearing loss^25^), then this explanation predicts that hearing loss at any age should degrade spatial attention, so older adults with age-related hearing loss should have weaker spatial attention than (older and younger) adults with normal hearing.

On the other hand, spatial attention may not scale with spatial acuity. In the aforementioned studies of auditory attention, stimuli were separated by much greater angles (i.e., greater than 15 degrees) than the minimum audible angle. Thus, reduced spatial acuity may not affect spatial attention at these angles of spatial separation. Also, even if *early-onset* hearing loss weakens spatial attention, spatial attention could be preserved in older adults with age-related hearing loss because they have considerable experience directing attention to sounds before they acquired hearing loss. Therefore, older adults might deploy spatial attention to a similar extent as adults with normal hearing—or perhaps even to a greater extent to help compensate for sensory decline.

Here, we compared the extent to which older and younger adults use centrally-presented visual spatial cues to improve speech intelligibility in a cocktail-party listening task. We examined whether preparatory spatial attention depends, in a graded manner, on the degree of hearing loss and spatial resolution—to separate peripheral–cognitive interactions from changes to attention that occur with age independently of peripheral function. Given that speech intelligibility in noisy environments has been hypothesised to relate to overall cognitive ability^26,27^, we also included a visual matrix reasoning test (often used to assess fluid intelligence^28^), to examine whether performance relates to the benefit that each participant gains from orienting endogenous spatial attention.

## Results

### Older and younger adults gain a similar benefit to the accuracy of speech intelligibility from advance spatial cues

We measured accuracy and reaction times (RTs) in a spatialised cocktail party listening paradigm (Fig. 1a), in which three talkers spoke different sentences simultaneously from different locations. Participants were asked to report the sentence spoken by a talker at a particular location, which was indicated by a visual spatial cue that was presented either 2000 or 100 ms before the acoustic stimuli began. Responses were recorded as correct if both the colour and number words were correctly identified from the target sentence. We calculated the proportion of correct responses at each TMR, separately for the longer and shorter cue-target interval conditions, and separately for each group.

**Fig. 1.**
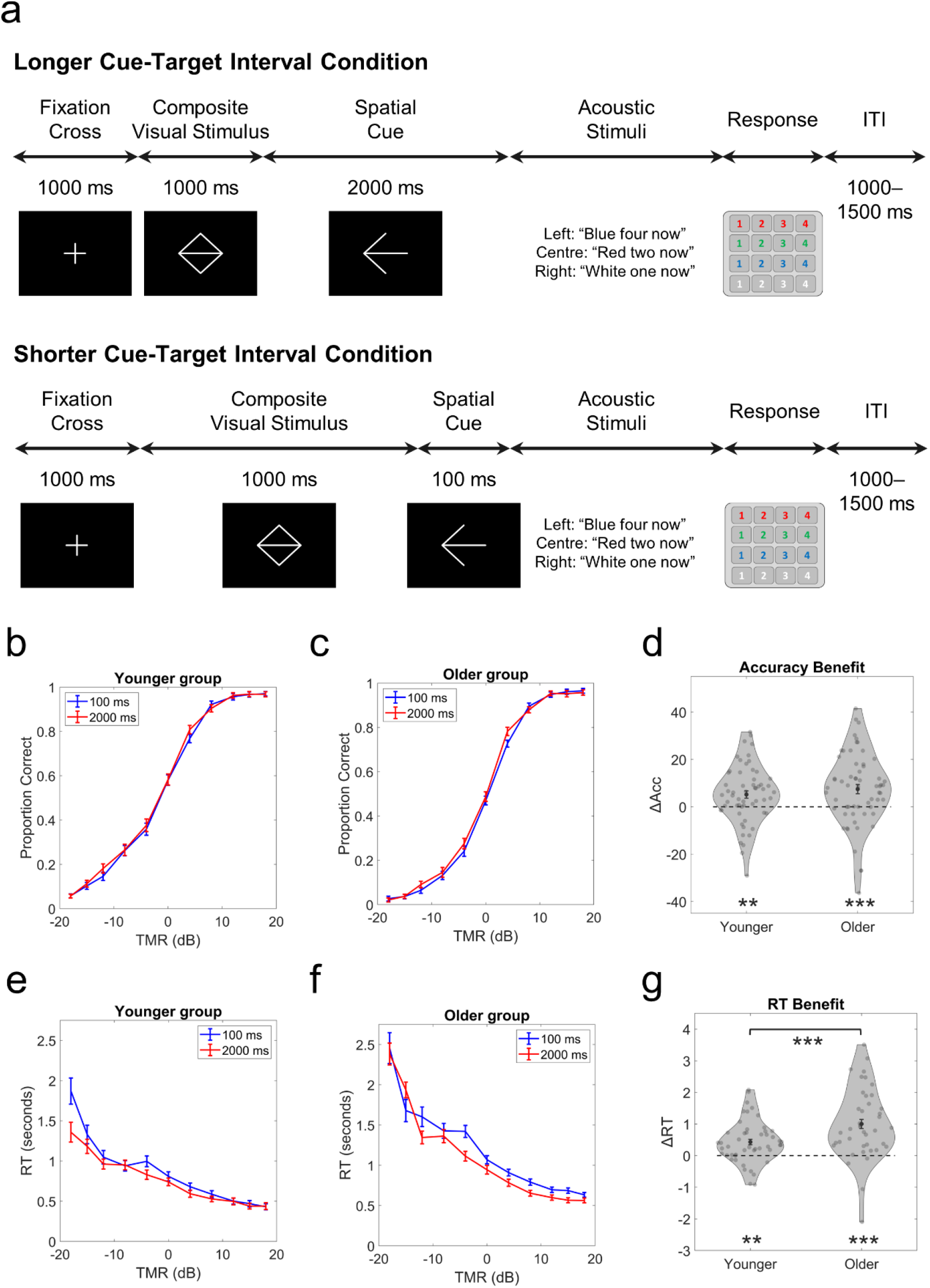
Experimental design and performance across groups. (a) Schematic of the trial structure for the longer and shorter cue-target interval conditions. The visual cue was an arrow that either pointed to the left (as depicted) or to the right. The three acoustic phrases were spoken by different talkers and simulated to originate from left, central, and right locations using interaural time differences (ITDs). ITI: Inter-trial interval. (b) Proportion of correct responses in the spatial attention task for the younger group, on average, at each cue-target interval condition (100, 2000 ms) and each target-to-masker ratio (TMR) condition. (c) Proportion of correct responses in the spatial attention task for the older group. (d) The benefit to accuracy from preparatory spatial attention, calculated as the difference in accuracy between the longer and shorter cue-target interval conditions. Black dots and black bars indicate the means and standard errors in each group, grey dots indicate results from individual participants, and the width of the violin plots indicates the distribution of results across participants. (e) Reaction times (RTs) measured from the onset of acoustic stimuli for the younger group, on average, across conditions. Error bars show ± 1 standard error of the mean. (f) RTs for the older group. (g) The benefit to RTs from preparatory spatial attention.

Fig. 1b–c illustrates the proportion correct across groups, TMRs, and cue-target intervals. As expected, accuracy improved with increasing TMR and was greater for the younger group (mean = .557, standard deviation = .104), overall, than for the older group (mean = .505, standard deviation = .077). There was also greater accuracy in the 2000-ms cue-target interval condition (mean = .535, standard deviation = .098) than the 100-ms cue-target interval condition (mean = .525, standard deviation = .093).

The three-way ANOVA revealed significant main effects of TMR, *F*(10, 1170) = 2038.3*, p* < .001*, ω_p_^2^* = .89, 95% CI = [.88, .90], Group, *F*(1, 117) = 9.60, *p* < .001*, ω_p_^2^* = .07, 95% CI = [.01, .18], and Cue-Target Interval, *F*(1, 1170) = 23.2, *p* < .001, *ω_p_^2^* < .01, 95% CI = [.00, .05].

The interaction between Group and TMR was significant, *F*(10, 1170) = 5.67, *p* < .001, *ω_p_^2^* = .02, 95% CI = [.00, .03], indicating that the difference in accuracy between the two age groups differed across TMRs. Follow-up independent-samples *t*-tests (with Bonferroni correction for multiple comparisons, ‘*p_bonf_’*) showed that the younger group had greater accuracy than the older group at -8 dB TMR [*t*(117) = 5.00, *p_bonf_* < .001, *d_s_* = .92, 95% CI = .54, 1.29], -4 dB TMR [*t*(117) = 4.23, *p_bonf_* = .007, *d_s_* = . 78, 95% CI = .40, 1.15], and 0 dB TMR [*t*(117) = 3.93, *p_bonf_* = .023, *d_s_* = .73, 95% CI = .35, 1.09], but there was no significant difference at the other TMRs, *t*(117) ≤ 3.51, *p_bonf_* ≥ .113, *d_s_* ≤ .65.

The interaction between Cue-Target Interval and TMR was significant, *F*(10, 1170) = 7.91, *p* < .001, *ω_p_^2^* = .01, 95% CI = [.00, .01], indicating that the difference in accuracy between the two cue-target interval conditions differed across TMRs. Follow-up paired-samples *t*-tests (with Bonferroni correction for multiple comparisons) showed that accuracy was greater in the longer cue-target interval condition than the shorter cue-target interval condition at -12 dB TMR [*t*(118) = 4.67, *p_bonf_* < .001, *d_z_* = .43, 95% CI = .24, .61], -4 dB TMR [*t*(118) = 3.92, *p_bonf_* = .021, *d_z_* = .36, 95% CI = .17, .54], and 4 dB TMR [*t*(118) = 7.20, *p_bonf_* < .001, *d_z_* = .66, 95% CI = .46, .86], but there was no significant difference at the other TMRs, *t*(118) ≤ 2.58, *p_bonf_* ∼ 1.00, *d_z_* ≤ .26.

We found no significant interaction between Cue-Target Interval and Group, *F*(1, 117) = .51, *p* = .477, *ω_p_^2^* < .01, 95% CI = [.00, 1.00], and no significant three-way interaction between Group, Cue-Target Interval, and TMR, *F*(10, 1170) = .59, *p* = .824, *ω_p_^2^* < .01, 95% CI = [.00, 1.00].

To compare the benefit from preparatory spatial attention between groups, we first calculated a metric to assess the benefit in each participant. For each participant in each cue-target interval condition, we fitted a sigmoid curve to proportion-correct scores across TMRs using a generalised linear model with a logistic link function (MATLAB function *glmfit* with argument ‘*logit’*). Next, we calculated the area under the curve for each cue-target interval condition. We took the difference in the area under the curve (ΔAcc) between the longer and shorter cue-target interval conditions as the benefit to speech intelligibility for each participant (with negative values indicating a decrement to speech intelligibility from the longer spatial cue). The advantage of using this metric (rather than comparing performance between cue-target interval conditions at individual TMRs) is that, by considering responses across a wide range of TMRs, we avoid ceiling and floor effects, and account for differences in overall speech intelligibility among individuals (and, therefore, between groups).

Fig. 1d illustrates the accuracy benefit from preparatory spatial attention (ΔAcc) for the two groups. One-sample *t*-tests showed a significant benefit (i.e., greater than zero) in both younger adults, *t*(57) = 3.23, *p* = .002, *d_z_* = .42, 95% CI = [.15, .69], and older adults, *t*(60) = 3.82, *p* < .001, *d_z_* = .49, 95% CI = [.22, .75]. An independent samples *t*-test showed that the magnitude of the benefit did not differ between groups, *t*(117) = .91, *p* = .367, *d_z_* = .17, 95% CI = [-.19, .53]. Given one of the main goals of this study was to compare the magnitude of the benefit from preparatory spatial attention between age groups, we followed up this non-significant difference by conducting a Bayesian equivalence test (using a Cauchy distribution centred around zero as a prior, which is the default prior in JASP). This allowed us to assess the evidence that the difference between groups is either identical or else has a small effect size (between -.10 and .10). The results showed a Bayes Factor of 3.50, which provides some evidence to support a similar magnitude benefit to speech intelligibility from preparatory spatial attention in younger and older adults.

### Older adults gain a greater benefit to reaction times from advance spatial cues than do younger adults

Fig. 1e–f illustrates RTs in the spatial attention task across Groups, TMRs, and Cue-Target Intervals. As expected, RTs were faster with increasing TMR, and the younger group had faster responses (mean = 1.68 seconds, standard deviation = .70) than the older group (mean = 2.06 seconds, standard deviation = .34). RTs were faster in the 2000-ms cue-target interval condition (mean = 1.81 seconds, standard deviation = .34) than the 100-ms cue-target interval condition (mean = 1.91 seconds, standard deviation = .39).

All three main effects were significant [TMR: *F*(7, 609) = 180.7, *p* < .001, *ω_p_^2^* = .31, 95% CI = .24, .36; Group: *F*(1, 87) = 9.36, *p* = .003, *ω_p_^2^* = .07, 95% CI = .00, .19; Cue-Target Interval: *F*(1, 87) = 40.6, *p <* .001, *ω_p_^2^* = .02, 95% CI = .00, .11]. Although, these main effects were qualified by significant two-way interactions.

The interaction between Group and TMR was significant, *F*(7, 609) = 4.53, *p <* .001, *ω_p_^2^* = .01, 95% CI = [.00, .02], showing that age group had different effects on RTs depending on the TMR. Follow-up independent samples *t*-tests (with Bonferroni correction for multiple comparisons) indicated that this interaction was driven by slower RTs in the older group compared to the younger group at -8 dB TMR, *t*(94) = 4.63, *p_bonf_* < .001, *d_s_* = .96, 95% CI = [.52, 1.37], and at -4 dB TMR, *t*(105) = 4.18, *p_bonf_* = .006, *d_s_* = .82, 95% CI = [.41, 1.20]; whereas, there were no significant differences in RTs between groups at the other TMRs, *t*(≥ 85) ≤ 2.41, *p_bonf_* ∼ 1.00, *d_s_* ≤ .47.

A significant interaction between Cue-Target Interval and TMR, *F*(7, 609) = 5.03, *p* < .001, *ω_p_^2^* = .01, 95% CI = [.00, .01], the difference in RTs between the two cue-target interval conditions differed across TMRs. Paired-sample t-tests (with Bonferroni correction for multiple comparisons) showed faster RTs in the longer cue-target interval condition than the shorter cue-target interval condition at -4 dB TMR [*t*(106) = 7.35, *p_bonf_* < .001, *d_z_* = .71, 95% CI = .50, .92] and 4 dB TMR [*t*(108) = 3.84, *p_bonf_* = .016, *d_z_* = .37, 95% CI = .17, .56], but no significant difference at the other TMRs, *t*(88) ≤ 3.39, *p_bonf_* ≥ .090, *d_z_* ≤ .27.

We also found a significant interaction between Group and Cue-Target Interval, *F*(1, 87) = 6.91, *p* = .010, *ω_p_^2^* < .01, 95% CI = [.00, .07]. Follow-up independent samples *t*-tests (with Bonferroni correction for multiple comparisons) showed a significant difference in RTs between groups at the shorter cue-target interval, *t*(87) = 3.55, *p_bonf_* = .004, *d_z_* = .76, 95% CI [.32, 1.18], but no significant difference at the longer cue-target interval, *t*(87) = 2.43, *p_bonf_* = .103, *d_z_* = .52, 95% CI = [.09, .94].

The three-way Group x Cue-Target Interval x TMR interaction was not significant, *F*(7, 609) = .412, *p* = .895, *ω_p_^2^* < .01, 95% CI [.00, 1.00].

Similar to accuracy, we calculated a metric to assess the benefit to RTs from preparatory spatial attention across TMRs, so we could compare the benefit to RTs between groups. For each participant in each cue-target interval condition, we fit a polygonal line to the RT data across TMRs from -8 dB to +18 dB TMR. We then calculated the difference (ΔRT) between the fitted lines for the longer and shorter cue-target interval conditions for each participant, which we took as the RT benefit (with negative values indicating a decrement to—i.e., an increase in—RTs from the longer spatial cue).

Fig. 1g illustrates the RT benefit from preparatory spatial attention (ΔRT) for the two groups. One-sample *t*-tests showed a significant benefit in both younger adults, *t*(57) = 4.74, *p* = .002, *d_z_* = .62,, 95% CI = [.34, .90] and older adults *t*(60) = 7.49, *p* < .001, *d_z_* = .96, 95% CI = [.65, 1.26]. An independent *t*-test revealed that the RT benefit was significantly larger in the older group than in the younger group, t(117) = 3.62, p < .001, *d_z_* = .67, 95% CI = [.29, 1.03].

### Individual differences in task performance relate to spatial acuity, matrix reasoning scores, and hearing sensitivity

Given we were interested in individual differences among participants, we conducted several analyses to identify relationships among tasks. To examine how performance relates to hearing loss, we included pure-tone audiometric thresholds at 4–8 kHz, averaged across the two ears (hereafter, ‘Audiogram’), This measure of high-frequency hearing captures the typical changes in older adults (for more detail, see Methods). We also included ITD thresholds (i.e., spatial discrimination thresholds corresponding to the minimal audible angle), and age-scaled matrix reasoning scores^29^ (hereafter, ‘WAIS-MR’). For these analyses, we examined relationships among variables in the younger and older groups separately.

First, given that most previous studies examining relationships among variables have used raw speech intelligibility scores^24,26^ (rather than the benefit from a longer cue-target interval), we investigated how individual differences in overall accuracy and RTs (collapsed across TMRs) on the spatial attention task relates to the other variables. For each group, we examined correlations between 6 pairs of variables (Audiogram, ITD Threshold, and WAIS-MR score with overall Accuracy and with RTs in the spatial attention task). The significance threshold was adjusted for multiple comparisons using the Bonferroni correction (p < .05 / 6 = .0083).

For the younger group, better (i.e., smaller) ITD Thresholds were associated with better Accuracy in the spatial attention task (*ρ* = -.49, *p* < .001; Fig. 2a). In addition, better WAIS-MR scores were associated with better Accuracy (*ρ* = .43, *p* < .001; Fig. 2a). None of the other correlations with Accuracy or RTs were significant (see Table 1). To test whether WAIS-MR and ITD Threshold explained separate variance in Accuracy in the younger group, we entered these variables into a stepwise linear regression. The results showed that WAIS-MR and ITD Threshold made significant independent contributions to predicting Accuracy in the spatial attention task (model containing ITD Threshold only: *R^2^* = .37, *p* < .001; model containing ITD Threshold and WAIS-MR: *R^2^* = .46, *R^2^* change = .09, *p* change = .005).

**Fig. 2.**
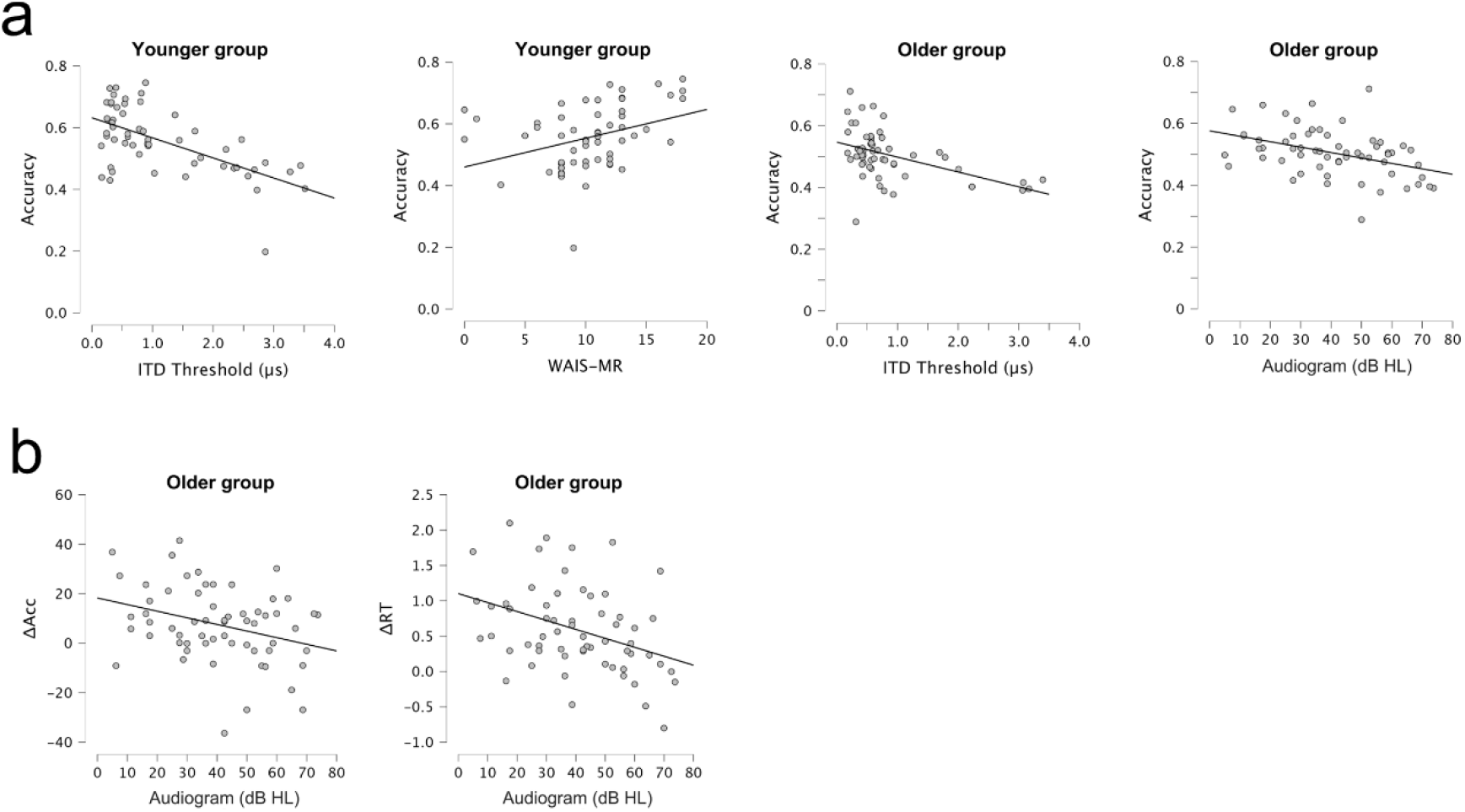
Scatter plots showing significant relationships between variables. (a) Significant relationships with overall accuracy in the spatial attention task: ITD Thresholds and WAIS-MR age-scaled scores for the younger group; ITD Thresholds and the Audiogram for the younger group. Individual points each represent one participant, and the solid lines display the lines of best fit. (b) Significant relationships between the Audiogram and the benefit from the longer cue-target interval in the older group, both for the benefit to accuracy (ΔAcc) and the benefit to reaction times (ΔRT). Positive values indicate better performance (i.e., greater accuracy or faster RTs in the longer cue-target interval condition).

**Table 1.** Spearman’s rho (*ρ*) and associated *p*-values for relationships with overall accuracy and reaction times (RTs) in the spatial attention task. Relationships that surpass the Bonferroni-corrected significance threshold (p < .0083) are indicated with bold font.

| Variable 1 | Variable 2 | Younger group |  | Older group |  |
| --- | --- | --- | --- | --- | --- |
| | | $\rho$ | $p$ | $\rho$ | $p$ |
| Audiogram | Accuracy | -.14 | .283 | **-.43\*** | **< .001\*** |
| ITD Threshold | Accuracy | **-.49\*** | **< .001\*** | **-.49\*** | **< .001\*** |
| WAIS-MR | Accuracy | **.43\*** | **< .001\*** | .30 | .020 |
| Audiogram | RTs | .13 | .319 | .26 | .044 |
| ITD Threshold | RTs | .28 | .033 | .06 | .683 |
| WAIS-MR | RTs | .02 | .870 | .32 | .013 |

For the older group, we found a significant correlation between ITD Thresholds and Accuracy, *ρ* = -.49, *p* < .001 (Fig. 2a), like for the younger group. We also found a significant correlation between the Audiogram and Accuracy, *ρ* = -.43, *p* < .001 (Fig. 2a). None of the other correlations were significant (see Table 1). Including both ITD Threshold and the Audiogram in a stepwise linear regression showed that these two variables made significant independent contributions to predicting Accuracy in the older group (model containing ITD Threshold only: *R^2^* = .23, *p* < .001; model containing ITD Threshold and Audiogram: *R^2^* = .29, *R^2^* change = .06, *p* change = .037).

### Individual differences in orienting auditory spatial attention among older adults are predicted by hearing sensitivity

While the previous analyses examined individual differences in overall performance, we were primarily interested in how the benefit from preparatory spatial attention varied across participants. We, therefore, examined relationships between the benefit to accuracy (ΔAcc) and benefit to RTs (ΔRT) from the longer cue-target interval with the Audiogram, ITD Threshold, and WAIS-MR score.

In the younger group, none of the correlations were significant, for either the benefit to accuracy or to RTs (Table 2).

**Table 2.** Spearman’s rho (*ρ*) and associated *p*-values for relationships with the benefit to accuracy (ΔAcc) and to reaction times (ΔRT) from preparatory spatial attention. Relationships that surpass the significance threshold (*p* < .05) are indicated with bold font.

| *Variable 1* | *Variable 2* | *Younger group* |  | *Older group* |  |
| --- | --- | --- | --- | --- | --- |
| | | $\rho$ | $p$ | $\rho$ | $p$ |
| Audiogram | $\Delta\text{Acc}$ | -.11 | .419 | **-.27** | **.037** |
| ITD Threshold | $\Delta\text{Acc}$ | -.25 | .067 | -.24 | .065 |
| WAIS-MR | $\Delta\text{Acc}$ | -.18 | .184 | .04 | .777 |
| Audiogram | $\Delta\text{RT}$ | -.03 | .849 | **-.39** | **.002** |
| ITD Threshold | $\Delta\text{RT}$ | -.10 | .462 | .12 | .357 |
| WAIS-MR | $\Delta\text{RT}$ | -.20 | .136 | -.14 | .294 |

In the older group, there were no significant correlations with ITD Threshold or WAIS-MR scores (Table 2). However, the Audiogram correlated significantly with both the benefit to accuracy (*ρ* = -.27, *p* = .037; Fig. 2b) and the benefit to RTs (*ρ* = .39, *p* = .002; Fig. 2b). The direction of these correlations indicates that greater hearing loss reduces the benefit from preparatory spatial attention, both when measured in terms of the improvement in accuracy and the improvement in RTs. We did not correct for multiple comparisons in this analysis, because these were planned comparisons related to the main hypothesis. Although, it is interesting to note that the correlation with ΔRT would remain significant after Bonferroni correction. To assess whether this relationship was a by-product of older age, we ran a follow-up partial Spearman’s correlation controlling for age in years. The correlation between the Audiogram and ΔRT remained significant after controlling for age (*ρ* = .44, *p* = .001).

Given that the younger and older groups showed different relationships between the Audiogram and the benefit from preparatory spatial attention, we conducted follow-up (two-tailed) bootstrapping analyses to compare the Spearman’s correlation coefficients between the two groups. The correlation between the Audiogram and ΔRT was significantly greater in older than younger adults [observed Δ*⍴* = .40, *p* = .002; null distribution mean Δ*⍴* < .01, 95% CI = -.27, .26]. Although, there was no significant difference in correlation coefficients between the two groups for the relationship between the Audiogram and ΔAcc [observed Δ*⍴* = .19, *p* = .083; null distribution mean Δ*⍴* < .001, 95% CI = -.25, .26].

## Discussion

We sought to disentangle how spatial attention relates to ageing, in general, and to sensory declines that occur with older age. We found the benefit to speech perception from preparatory spatial attention is at least as large in older as in younger adults, as a group (Fig. 1d). Thus, consistent with previous work on visual endogenous attention^2–4^, people aged 55 and above generally retain the capacity to orient auditory spatial attention. Nevertheless, our results demonstrate that sensory–cognitive interactions, which have not been explicitly modelled in previous studies, are crucial for understanding attention in older age. We found that sensory processing affected the magnitude of the benefit from preparatory spatial attention: Greater age-related hearing loss progressively reduced the benefit (Fig. 2b). Thus, while spatial selective auditory attention is preserved in older age, it is degraded by age-related hearing loss. Given that more than 40% of adults aged 55 and above experience declines in hearing sensitivity^10^, our findings highlight the importance of considering peripheral sensory processing as a crucial predictor of cognitive abilities in older age.

Our results extend previous results showing impaired preparatory spatial attention in children with early-onset hearing loss^19^, by showing this also applies to hearing loss occurring much later in the lifespan. In addition, our results indicate that the ability to orient spatial attention is not all-or-none but rather varies in magnitude across individuals depending on the degree of hearing loss. In addition to the difficulties understanding speech posed by reduced audibility, declines in spatial attention may make it more difficult for individuals to perceptually segregate talkers in situations with multiple competing talkers (which is supported by different patterns of errors between older and younger adults: see Supplemental File). Our finding is particularly noteworthy because most of our sample have audiometric thresholds that would not lead to a clinical diagnosis of hearing loss. Thus, even small variations in hearing sensitivity impact endogenous spatial attention, which could help to explain why many middle-aged adults report difficulties understanding speech in noisy environments, despite not meeting the criteria for a hearing-loss diagnosis^30,31^.

In general, our findings oppose a compensatory mechanism for orienting spatial attention related to deteriorations in sensory processing, because under this explanation we would expect a *greater* benefit from preparatory spatial attention in people who had more age-related hearing loss, whereas our results showed the opposite pattern. Instead, our results support a more nuanced explanation that may involve multiple mechanisms. The first mechanism relates to the enhancement of the RT benefit from preparatory spatial attention in older age, which is unrelated to sensory processing and could instead compensate for other processes that change with age— such as processing speed^32–34^ and/or response speed^35,36^. Alternatively, we might have only found a greater benefit to RTs because baseline RTs (in the shorter cue-target interval condition) were slower in older adults and, therefore, their RTs had greater room to improve. Under the latter argument, no compensatory mechanism would be required to explain our results.

A second mechanism is needed to account for the finding that declines in sensory processing *reduce* the benefit from preparatory spatial attention: Possibly, people with hearing loss are unable to benefit from preparatory spatial attention or, alternatively, they may have learned to diminish orienting spatial attention as a listening ‘strategy.’ Interestingly, although overall speech intelligibility in our spatial attention task depended on a listener’s spatial acuity (consistent with Dai *et al.*^24^), we found no evidence that the magnitude of the benefit from preparatory spatial attention was related to spatial acuity. This implies that the mechanism by which age-related hearing loss affects preparatory spatial attention is not due to reduced spatial acuity. The exact mechanism is unclear, although could possibly be underpinned by broader changes in the ‘strategies’ that people use to accommodate changes in hearing; for example, reduced audibility may encourage greater reliance on semantic context^37,38^ and decrease reliance on other (e.g., spatial) information. Alternatively, hearing loss may place greater cognitive demand on understanding speech in noisy environments^39^, and—despite benefiting speech intelligibility in the long term—deploying preparatory spatial attention could require additional cognitive resources that are less available under greater cognitive demand.

Consistent with previous research, we found that overall speech intelligibility (across all of the cue-target interval and TMR conditions we tested) was worse for older listeners who had poorer hearing sensitivity^40^ (Fig. 2a) or poorer spatial acuity^24^ (Fig. 2a). The relationship with spatial acuity persisted in both older and younger adults, implying that even small differences in spatial acuity among young adults without hearing loss can help to predict their ability to understand speech in spatialised settings. Finally, WAIS Matrix Reasoning scores covaried with overall performance (Fig. 2a; Supplemental Fig. 2). We assume this relationship is likely related to fluid intelligence, rather than to the spatial component of the matrix reasoning task, given that a previous study found a relationship between matrix reasoning and speech intelligibility in a non-spatialised sentence-in-noise task^41^.

Our findings may have implications for understanding links between hearing loss and dementia^42^. We studied healthy older adults with no known cognitive impairments using a cross-sectional design, which precludes inferences about causality or directionality. Nevertheless, our results provide an example of how one specific cognitive function—auditory spatial orienting—is linked to subtle changes in peripheral hearing (not mediated through changes in spatial acuity). Effects on preparatory spatial attention are relevant to real-world listening and were not considered in Griffiths *et al.*^8^. Griffiths *et al.*^8^ did present listening effort as a critical determinant of dementia linked to hearing loss, separate from hearing sensitivity, and suggested that measures of speech-in-noise might be better predictors of dementia than the pure-tone audiogram. However, the current results suggest there could be value in assessing attention during spatialised listening in addition to other measures, which could be accomplished using the type of spatial attention test we used here.

In conclusion, while endogenous auditory spatial orienting is generally preserved in older age, it is degraded by age-related hearing loss—demonstrating sensory– cognitive interactions in ageing. Interestingly, the degradation of auditory spatial attention is not explained by poorer spatial acuity associated with hearing loss. Given that sensory decline is extremely prevalent among older adults, these sensory– cognitive interactions are more likely to be the norm than the exception, and are necessary for predicting real-world perception among older populations.

## Methods

### Participants

We recruited 61 younger adults aged 18–35 years and 64 older adults aged 55–81 years. Of these, we excluded six participants (three from each group): one younger participant had recent ear surgery, two (1 older, 1 younger) withdrew part-way through the study, two (1 older, 1 younger) did not follow the task instructions (after the experiment, they told us that they ignored the attentional cues during the spatial attention task), and one older participant had below-chance accuracy across all conditions of the spatial attention task. Of the remaining participants, two older participants completed most (80–90%) of the main task and were included in the analyses. Thus, we analysed the data from 58 younger (25 male, 33 female; median age = 24 years, interquartile range [IQR] = 8) and 61 older (22 male, 39 female; median age = 70 years, IQR = 7) participants. This sample size is sensitive to correlations (e.g., between performance metrics and audiometric thresholds; N=119; power = .8) of size *r*^2^ ≥ .06, and differences in Pearson’s correlation coefficient (i.e. different relationships between performance and audiometric thresholds in older and younger groups) of size Cohen’s *q* ≥ .5.

All participants were native English speakers and had normal or corrected-to-normal vision. We measured their pure-tone threshold at both ears (at octave frequencies between 250 and 8000 Hz) average pure-tone thresholds. Pure-tone thresholds were measured in accordance with BS EN ISO 8253-1^43^ using a Starkey Acoustic Analyser AA30 (Starkey Laboratories, Inc.). Audiograms for all participants are illustrated in Fig. 3. The younger group had average pure-tone thresholds of 8.32 dB HL (range = 0 – 47.5) and the older group had average pure-tone thresholds of 14.06 dB HL (range = -5 – 30). For all participants, the difference in six-frequency average pure-tone thresholds between the left and right ears was 22.5 dB or less (mean = 6.52 dB, standard deviation = 3.76). Participants who used hearing aids (N = 10, all in the older group) removed them while completing the tasks.

**Fig. 3.**
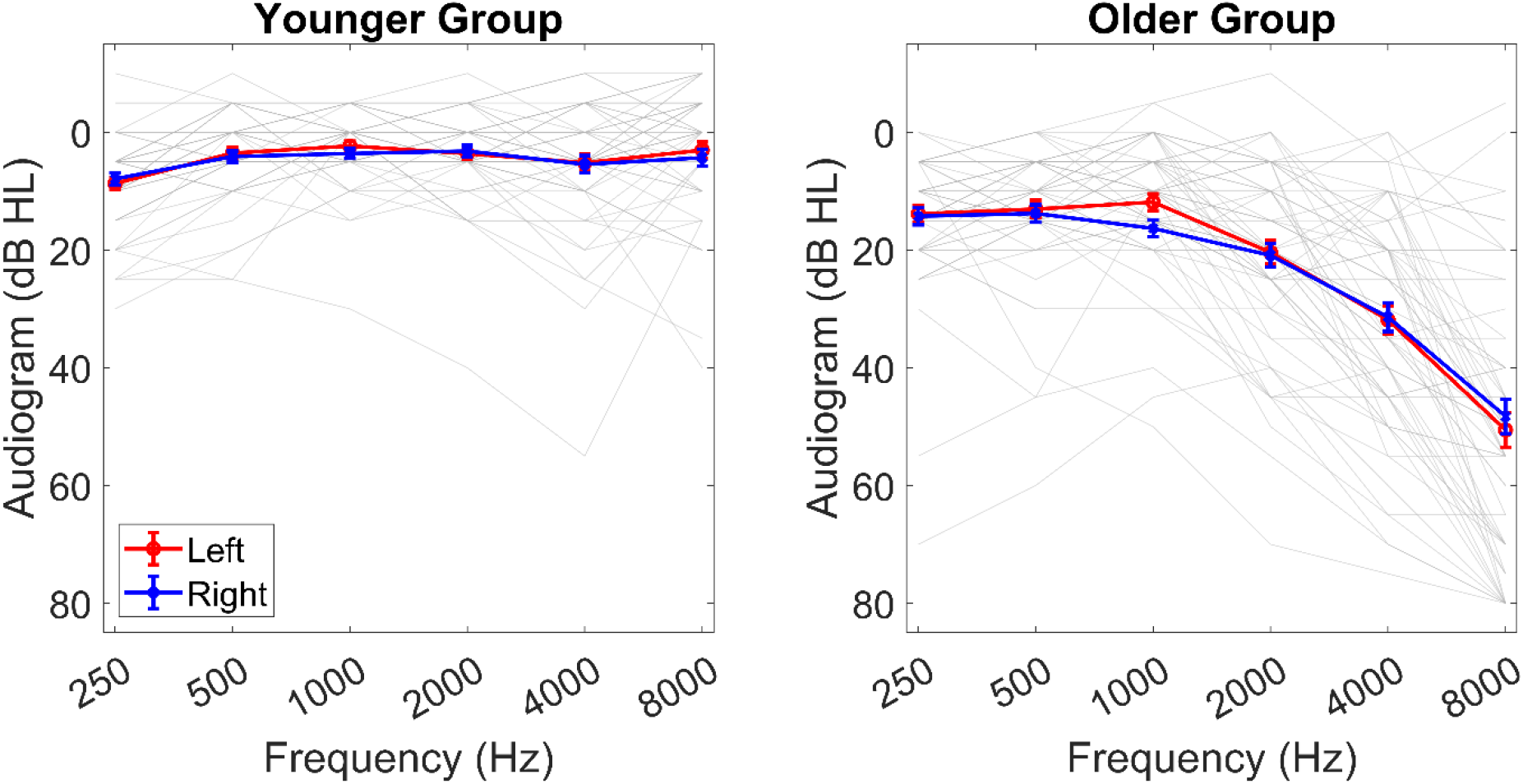
Pure-tone audiometric thresholds for the younger and older groups. Blue and red lines indicate the averages for the right and left ears in each group. Grey lines indicate the averages across both ears for individual participants.

The study was approved by the UCL Research Ethics Committee. All participants gave informed consent and were compensated for their time at standard UCL rates.

## Materials and Procedures

The experiment was conducted in a sound-attenuating booth. Participants sat in a chair facing an LCD visual display unit (Dell Inc.). All tasks were delivered using custom-written MATLAB scripts (version R2019a; MathWorks, Inc.) with Psychtoolbox^44^ (version: 3.0.17). Acoustic stimuli were presented through an external sound card (ESI Maya 22 USB; ESI Audiotechnik GmbH, Leonberg) connected to circumaural headphones (Sennheiser HD 380 Pro; Sennheiser electronic GmbH & Co. KG). Participants’ responses were recorded using a touch screen (iiyama ProLite T2435MSC; iiyama Corporation).

### WAIS matrix reasoning task

Participants first completed a computerised version of the Wechsler Adult Intelligence Scale-IV (WAIS-IV) Matrix Reasoning test^29^. On each trial, participants viewed an array of images, which contained one missing image. They were presented with five options and were instructed to select the image that fitted into the array. Participants were provided with two examples with written explanations before commencing the test.

### Spatial attention task

Next, participants completed a spatial attention task. Acoustic stimuli were modified phrases from the Co-ordinate Response Measure corpus^45^ (CRM) used by Holmes *et al.*^13,19^. Each phrase contained a colour word (“Red”, “Green”, “Blue”, or “White”), a number word (“one”, “two”, “three”, or “four”), and the word “now”; for example, “Red two now”. The phrases were spoken by native British English talkers. We wanted to present three distinct voices on each trial; given that age and gender are salient voice cues, we used one male voice, one female voice, and one “child” voice, which included phrases spoken by a different female talker that were manipulated using Praat^©^ (Version 5.3.08; http://www.praat.org/) to have a higher fundamental frequency and shorter formant spacing ratio than the original recordings. The digital recordings had an average duration of 1.4 seconds and were all adjusted to have the same root mean square (RMS) power. Visual stimuli were white arrows on a black background that pointed leftwards or rightwards (left and right spatial cues, respectively), and a composite stimulus that contained both arrows overlaid.

Fig. 1a illustrates the trial structure for the spatial attention task. On every trial, participants saw the visual composite stimulus (which was uninformative about the target location) followed by the left or right spatial cue, which indicated the location of the target talker and varied from trial to trial. There were two cue-target interval conditions: In the longer cue-target interval condition, the composite stimulus was presented for 1000 ms, then the spatial cue was presented for 2000 ms before the acoustic stimuli began; in the shorter cue-target interval condition, the composite stimulus was presented for 2900 ms, then the spatial cue was presented for 100 ms before the acoustic stimuli began. In all conditions, the spatial cue remained on the screen until the acoustic stimuli had ended, so participants did not have to maintain the cue in memory. The acoustic stimuli for each trial consisted of three overlaid phrases, which were always spoken by different talkers, contained different colour and number words, and were simulated to come from different spatial locations using ITDs. The child’s voice was always presented with an ITD of 0 µs; in other words, it was simulated to come from a central location and was never the target. The locations of the male and female voices varied pseudo-randomly on each trial. One had an ITD of -205 µs, and was therefore simulated to the left side. The other had an ITD of +205 µs, and was therefore simulated to the right side. For each trial, the target-to-masker ratio (TMR) was selected from one of eleven values (-18, -15, -12, -8, -4, 0, +4, +8, +12, +15, or +18 dB), which specified the level difference between the target phrase and each of the two competing phrases. The overall amplitude of the combined phrases was set to the same RMS value across trials. Participants were instructed to listen to the voice that corresponded to the direction of the arrow, and to report the colour-number combination from the target phrase by pressing one of sixteen coloured digits on the touch screen, which were displayed on the touch screen from the beginning of each trial. Participants were instructed to respond as quickly and as accurately as possible, and to guess if uncertain. The intertrial interval was randomly selected on each trial (1.0–1.5 seconds).

Participants first completed 12 practice trials, which were identical to the main task except that explicit feedback was provided (i.e., “correct” or “incorrect” was displayed on the screen after each trial). They then completed the main task, which consisted of 440 trials without feedback, separated into 10 blocks of 44 trials. Each block contained one trial from each condition (2 cue-target intervals x 2 spatial cue directions x 11 TMRs).

### ITD discrimination task

After the spatial attention task, participants completed an adaptive ITD discrimination task using a procedure similar to Dai *et al.*^24^, except that the stimuli were spoken phrases from the spatial attention task rather than complex tones. On each trial, participants heard two phrases spoken one after another. The second phrase was spoken by the “child” and was located centrally (ITD = 0 µs), while the first sentence was spoken by the male or female talker and was either to the left (negative ITD) or right (positive ITD). Participants were asked to report whether the first sentence was to the left or the right by pressing buttons on the touch screen that were labelled “left” and “right”. The buttons were displayed from the beginning of each trial. Participants were instructed to respond as accurately as possible, and told that they did not need to respond quickly. The ITD was manipulated using an 1-up 3-down adaptive procedure, with a starting ITD of 287 µs and an initial step size of 12.8 µs, which decreased to .512 µs after 3 reversals. The run ended after the participant reached a maximum of 10 reversals, reached the minimum ITD value of .28 µs, or exceeded 192 trials (whichever happened first). The ITD was adapted in separate runs for the male and female voice, and the order of the two runs was counterbalanced across participants.

For each run, we calculated the ITD threshold as the median of the final 6 reversals, then took the average across the two runs for each participant. Three participants reached the minimum ITD value for one run and, for these participants, we used a value of 0 as their ‘threshold’ for that run. Three participants reached the maximum number of trials on both runs, so we excluded them from further analyses involving the ITD threshold (although, we included these participants in other analyses not involving ITD thresholds). For participants who reached the maximum number of trials on either the male (N = 13) or the female (N = 8) run, we used the other run as their ITD threshold.

## Analyses

Statistical analyses were conducted with MATLAB^©^ (version R2022a, The MathWorks, Inc., Natick, MA, USA) and JASP™ (versions 0.17.2.1^46^ and 0.18.1^47^). We calculated effect sizes and confidence intervals (CIs) using MOTE^48^. All tests are two-tailed.

### Spatial attention task: Accuracy

We used a three-way mixed ANOVA to compare the effects of Group (younger or older; between-subjects variable), TMR (11 levels from -18 to +18 dB; within-subjects variable), and Cue-Target Interval (2000 or 100 ms; within-subjects variable) on the proportion of correct responses.

We compared the benefit from the longer cue-target interval (ΔAcc) between groups using an independent samples *t*-test.

### Spatial attention task: Reaction Times

Reaction times (RTs) were measured from the onset of the acoustic stimuli. We analysed RTs for correct trials only, and only at TMRs from -8 dB to +18 dB, because many participants had no correct trials below -8 dB TMR. The cut-off TMR of -8 dB was chosen to ensure that at least 70% of participants in both groups had at least one correct trial in each Cue-Target Interval condition (for comparison, 100% of participants had at least one correct trial per condition at all positive TMRs, but below -8 dB TMR, ≤ 51% of participants in the older group had at least one correct trial per condition). We used a three-way mixed ANOVA to compare the effects of Group, TMR, and Cue-Target Interval on RTs.

We compared the RT benefit from the longer cue-target interval (ΔRT) between groups using an independent samples *t*-test.

### Individual differences: Relationships among tasks

We used Spearman’s correlation coefficients to examine the relationships among tasks, due to non-normality of residuals and heteroskedasticity in the data, and we used Bonferroni corrections for multiple comparisons. Where we obtained different relationships in the younger and older groups, we performed follow-up bootstrapping analyses (using 10,000 samples with replacement) to compare the Spearman’s correlation coefficients between groups, to test the hypothesis that the two coefficients are drawn from different distributions.

## Acknowledgments

This work was supported by a Pauline Ashley Fellowship from the Royal National Institute for Deaf People (RNID) to Emma Holmes (PA25_Holmes). This work was conducted at the Wellcome Centre for Human Neuroimaging, which is funded by Wellcome (Ref: 203147/Z/16/Z).

## Author contributions

Andrea Caso: Investigation, Formal analysis, Visualization, Writing – original draft. Timothy Griffiths: Conceptualization, Writing – editing. Emma Holmes: Conceptualization, Methodology, Software, Investigation, Formal analysis, Writing – original draft, Writing – editing, Supervision, Funding acquisition.

## Data availability statement

Experiment files, data, and analysis code for this study are publicly available on the Open Science Framework (OSF) and can be accessed at https://osf.io/gvys3/?view_only=f7f27642ef984c3aa10ab046135b8862.

