## Supplemental for "Spatial selective auditory attention is preserved in older age but is degraded by peripheral hearing loss"

### Supplemental Material

#### Error types for the spatial attention task

To identify how participants responded on incorrect trials, we categorised their responses into four different error types. A ‘masker error’ was recorded if they reported the colour and number words from the contralateral location. A ‘child error’ was recorded if they reported the colour and number words spoken by the child, which was always presented at the central location. A ‘mix error’ was recorded if the reported colour and number were both spoken on the current trial, but by two different talkers. Otherwise, one or both words were not spoken by any of the talkers and this was categorised as a ‘random error’. We calculated the proportion of each error type, out of the total number of incorrect trials, for each participant, collapsing across TMRs and cue-target interval conditions. We then conducted a two-way mixed ANOVA to compare the effects of Group (younger or older; between-subjects variable) and Error Type (masker, child, mixed, random; within-subjects variable) on the proportions of errors.

Supplemental Fig. 1 illustrates the proportions of the four error types across groups, collapsed across TMRs and cue-target intervals. Both groups showed the greatest proportion of Mix errors, followed by Masker errors, then Child errors, and then Random errors.

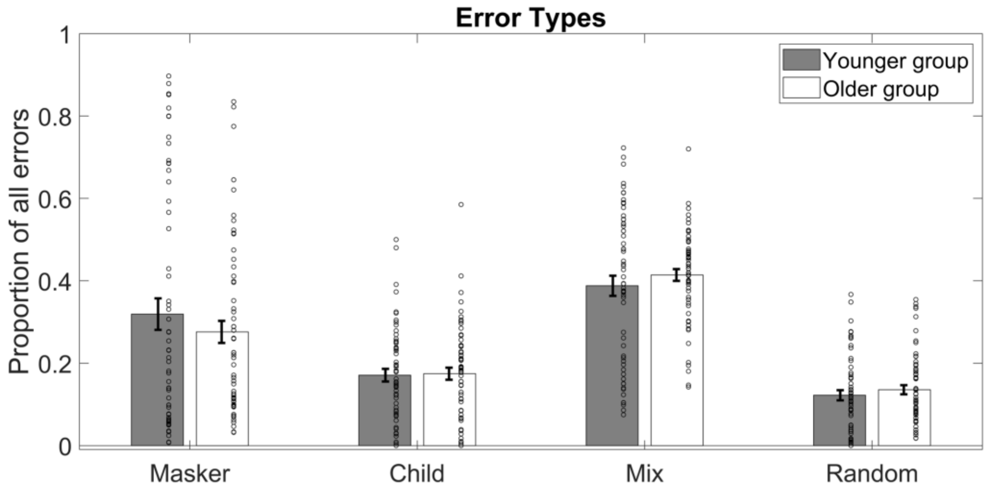


**Supplemental Fig. 1.** Bar graph showing proportions of each error type (as a proportion of all incorrect trials) for the younger and older groups. Error bars show ± 1 standard error of the mean.

The main effect of Error Type was significant, $F(3, 1170) = 45.5$, $p < .001$, $\omega_p^2 = .21$, 95% CI = [.13, .28]. Follow-up paired-samples *t*-tests (with Bonferroni correction for multiple comparisons) showed a significant difference between most pairs of Error Types, except from the proportion of Masker compared to Mix errors and the proportion of Child compared to Random errors [Masker and Child errors: $t(118) = 6.65$, $p_{bonf} < .001$, $d_z = .61$, 95% CI = .41, .80; Masker and Mix errors: $t(118) = -1.59$, $p = .068$, $d_z = -.15$, 95% CI = -.33, .04; Masker and Random errors: $t(118) = 8.01$, $p_{bonf} < .001$, $d_z = .73$, 95% CI = .53, .94; Child and Mix errors: $t(118) = -8.24$, $p_{bonf} < .001$, $d_z = -.76$, 95% CI = .55, .96; Child and Random errors: $t(118) = 1.36$, $p \sim 1.00$, $d_z = .12$, 95% CI = -.06, .30; Mix and Random errors: $t(118) = 9.60$, $p_{bonf} < .001$, $d_z = .88$, 95% CI = .67, 1.09].

The main effect of Group was not significant, $F(1, 117) = 3.44$, $p = .066$, $\omega_p^2 = .02$, 95% CI = [.00, .10]. Although, there was a significant interaction between Group and Error Type, $F(3, 1170) = 2.90$, $p = .035$, $\omega_p^2 < .01$, 95% CI = [.00 .03], which indicates that the types of errors participants made on incorrect trials differed between groups. Follow-up *t*-tests revealed that the interaction between Group and Error type was underpinned by a significant difference between Mix and Masker errors in the older group [$t(60) = 3.67$, $p_{bonf} = .002$, $d_z = .47$, 95% CI = .20, .73] that was not significant in the younger group [$t(57) = .85$, $p_{bonf} = .398$, $d_z = .11$, 95% CI = -.15, .37]. Whereas, differences between other pairs of error types showed the same pattern of significance in both groups [Masker and Child (Younger): $t(57) = 5.02$, $p_{bonf} < .001$, $d_z = .66$, 95% CI = .37, .94; Masker and Child (Older): $t(60) = 4.32$, $p_{bonf} = .026$, $d_z$ = ..55, 95% CI = .28, .82; Masker and Random (Younger): $t(57) = 5.83$, $p_{bonf} < .001$, $d_z = .77$, 95% CI = .47, 1.06; Masker and Random (Older): $t(60) = 5.48$, $p_{bonf} < .001$, $d_z = .70$, 95% CI = .42, .98; Child and Mix (Younger): $t(57) = 4.17$, $p_{bonf} < .001$, $d_z = .55$, 95% CI = .27, .82; Child and Mix (Older): $t(60) = 7.99$, $p_{bonf} < .001$, $d_z = 1.02$, 95% CI = .71, 1.33; Child and Random (Younger): $t(57) = .82$, $p_{bonf} \sim 1.00$, $d_z = .11$, 95% CI = -.15, .36; Child and Random (Older): $t(60) = 1.16$, $p_{bonf} \sim 1.00$, $d_z = .15$, 95% CI = -.10, .40; Mix and Random (Younger): $t(57) = 4.98$, $p_{bonf} < .001$, $d_z = .65$, 95% CI = .37, .94; Mix and Random (Older): $t(60) = 9.14$, $p_{bonf} < .001$, $d_z = 1.17$, 95% CI = .84, 1.49]. Given that Mix errors constitute responses that include words spoken by two different talkers, a greater proportion of Mix than Masker errors implies that, on incorrect trials, older adults experienced difficulty segregating the talkers from one another. In other words, when incorrect, the older group were more likely to erroneously group speech spoken by two different speakers than to segregate it.

#### LISAS for overall performance on the spatial attention task

There was no significant correlation between Accuracy and RTs in the spatial attention task (younger group: $\rho = -.20$, $p = .140$; older group: $\rho = -.25$, $p = .056$), so we decided to combine these two measurements into linear integrated speed-accuracy scores (LISAS[1]), to account for the idea that some participants may prioritise accuracy whereas others might prioritise RTs. We chose to do this because LISAS can accommodate individual differences in speed–accuracy trade-off[2] and has been used in past research on ageing[3,4]. Such integrated performance measures may be more sensitive to effects that are expected to be present in both speed and accuracy, especially when speed–accuracy trade-offs are present.

The LISAS is calculated by multiplying the proportion of errors by the standard deviation of RTs, divided by the standard deviation of the proportion of errors, with the goal of aligning RTs and accuracy into a comparable measurement scale before they are combined—such that the LISAS can be interpreted as an RT score that is corrected for the proportion of errors. The LISAS has the following analytical form (where j represents one participant, PE is the proportion of errors, RT is reaction time, and S represents the standard deviations for each variable):

$$LISAS_j = RT_j + \frac{S_{RT}}{S_{PE}} PE_j \quad \text{(Equation 1)}$$

For each group, we examined correlations between 3 pairs of variables (Audiogram, ITD Threshold, and WAIS-MR with LISAS), so the Bonferroni-corrected significance threshold was $p < .0167$.

For the younger group, we found a positive correlation between WAIS-MR scores and LISAS ($\rho = .39$, $p = .003$; Supplemental Fig. 2). Neither of the other correlations were significant (Audiogram and LISAS: $\rho = -.07$, $p = .602$; ITD Threshold and LISAS: $\rho = -.21$, $p = .114$). Removing one outlier who had a negative LISAS did not affect the pattern of results.

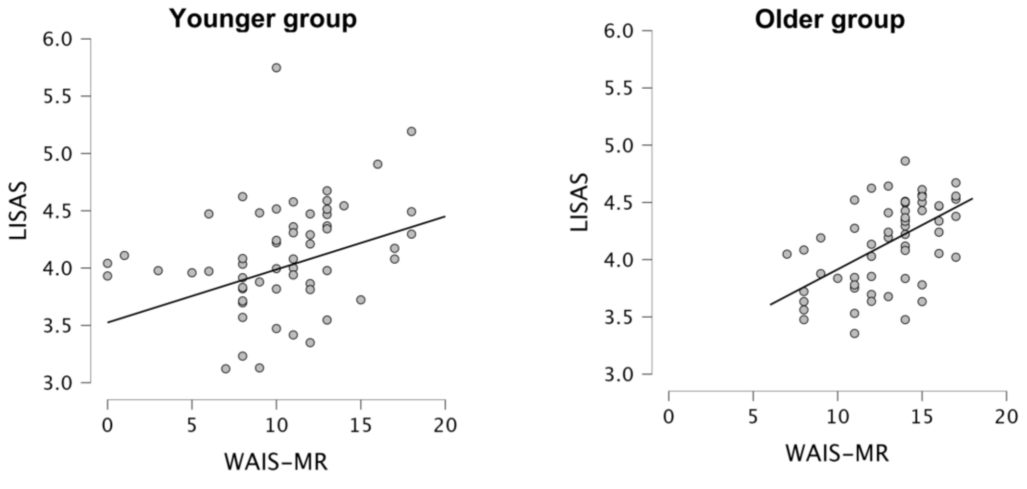


**Supplemental Fig. 2.** Significant relationships between WAIS-MR and LISAS in the spatial attention task for younger and older adults. Outliers (one in the younger group and one in the older group) have been excluded from these plots.

Similarly, in the older adult group, the only significant correlation was between WAIS-MR score and LISAS ($\rho = .47$, $p < .001$; Supplemental Fig. 2). Removing one outlier who scored zero in the WAIS-MR task increased the magnitude of the correlation coefficient ($\rho$ = .54, $p$ < .001). Thus, cognitive abilities correlate with integrated accuracy and RT scores in both groups. Neither of the other correlations met the Bonferroni-corrected significance threshold in the older group (Audiogram and LISAS: $\rho$ = -.09, $p$ = .493; ITD Threshold and LISAS: $\rho$ = -.27, $p$ = .039).

Based on these results, using LISAS was beneficial for examining individual differences in performance in the spatial attention task: We were able to detect a relationship between LISAS and WAIS Matrix Reasoning scores in the older group that was not apparent from either accuracy or RTs alone, but showed a consistent relationship across both groups when using LISAS as the dependent variable (Supplemental Fig. 2).

#### LISAS benefit from preparatory spatial attention

There was no significant correlation between the benefit to accuracy (ΔAcc) and the benefit to reaction times (ΔRT) from the longer cue-target interval in either group (younger group: $\rho$ = .163, $p$ = .221; older group: $\rho$ = -.001, $p$ = .993), so we ran some exploratory analyses with ΔLISAS, which integrates ΔAcc and ΔRT. We chose to do this in case some participants prioritise improvements in accuracy whereas others prioritise improvements in RT. However, we found no significant relationships between ΔLISAS and the variables of interest (Supplemental Table 1; Bonferroni-corrected significance threshold: $p < .05/3 = .0167$), suggesting that integrating the speed and accuracy benefits does not yield any additional insights in this instance. Thus, combining accuracy and RTs may be useful for examining individual differences in overall performance that account for a speed–accuracy trade-off (as examined in the section above), but may not be as useful for combining the *benefit* from preparatory spatial attention. A possible reason is that integrated measures of speed and accuracy (such as LISAS) assume that these two measures reflect overlapping cognitive processes, whereas the *benefits* of preparatory spatial attention to accuracy and RTs might be underpinned by distinct computational processes, as suggested by previous modelling work of preparatory spatial attention[5].

**Supplemental Table 1.** Spearman's rho ($\rho$) and associated $p$-values for relationships with the benefit to composite LISAS (ΔLISAS) from preparatory spatial attention. No relationships surpass the significance threshold.

| *Variable 1* | *Variable 2* | *Younger group* | | *Older group* | |
|---|---|---|---|---|---|
| | | $\rho$ | $p$ | $\rho$ | $p$ |
| Audiogram | ΔLISAS | .08 | .532 | -.08 | .540 |
| ITD Threshold | ΔLISAS | .23 | .086 | .25 | .063 |
| WAIS-MR | ΔLISAS | .26 | .053 | -.01 | .924 |
